# Origin and early divergence of tandem duplicated sorbitol transporter genes in Rosaceae: insights from evolutionary analysis of the SOT gene family in Angiosperms

**DOI:** 10.1101/2023.04.13.536778

**Authors:** Fan Yang, Jiawei Luo, Wenmeng Guo, Yuxin Zhang, Yunxiao Liu, Ze Yu, Yaqiang Sun, Mingjun Li, Fengwang Ma, Tao Zhao

## Abstract

Sorbitol is a critical photosynthate and storage substance in the Rosaceae family. Sorbitol transporters (SOTs) play a vital role in facilitating sorbitol allocation from source to sink organs and sugar accumulation in sink organs. Despite observing gene duplications in the SOT gene family, the origin and evolutionary process of these duplications are unclear, due to the complicated interplay of whole genome duplications and tandem duplications. Here, we investigated the synteny relationships of all detected Polyol/Monosaccharide Transporter (PLT) genes in 61 angiosperm genomes and SOT genes in major representative Rosaceae genomes. Combining phylogenetic analysis, we elucidated the lineage-specific expansion and syntenic conservation of PLTs and SOTs across different plant lineages. We found that Rosaceae SOTs, as PLT family members, originated from a pair of PLT tandem duplicating genes belonging to Class III-A. Additionally, lineage-specific and synergistic duplications in Amygdaloideae had contributed to the expansion of SOTs in Rosaceae plants. Overall, our findings provide insights into the genomic origins, duplication, and divergence of SOT gene family members, which are fundamental for further functional characterizations of each member.

## Introduction

Sugars are essential for all plant life activities, serving not only as a cellular energy source and structural component but also as signaling molecules that control plant growth and gene expression through complex signal transduction mechanisms (Stokes et al., 2013). Efficient and controlled distribution of sugar from source to sink is a crucial determinant of plant growth and development (Wang et al., 2013). While sucrose is the primary photosynthetic transport product of most fruit trees, synthesized in source tissues and transported to reservoir tissues through switched phloem-unloading symplasmic-to-apoplasmic pathway (Gu et al., 2021a, Slawinski et al., 2021), for Rosaceae species, the main photosynthate and soluble storage substance is sorbitol, which is distributed via sorbitol transporter (SOTs). SOTs are considered an important components of the Polyol/Monosaccharide Transporter (PMTs or PLTs) family. While few functional studies have focused on PLTs, numerous researchers have explored the roles of SOTs.

SOTs play a key role in long-distance of sugar transport, enabling sugar allocation from source to sink organs and the accumulation of sugars in sink organs (Zhang et al., 2014, Yu et al., 2019). In recent years, SOTs have been identified and characterized in many plant species. For example, in sour cherries, *PcSOT1* and *PcSOT2* were identified and shown to be highly expressed during fruit expansion and early stages of fruit development, respectively (Gao et al., 2003). Northern blot analysis demonstrated that *MdSOT3*/*4*/*5* in apple are related to sorbitol loading and accumulation in source organs (Watari et al., 2004). By analyzing the expression profiles of SOTs in various pear tissues, Yu et al identified 24 SOT genes from the pear genome and obtained the main effector gene *PbSOT6*/*20,* which was highly expressed in leaves, petioles, flowers, and fruit flesh (Yu et al., 2019). Recently, a domestication gene (MD10G1079800) affecting sorbitol accumulation was identified on apple chromosome 10, which encodes a sorbitol transporter (Gu et al., 2021b). The important SOT domestication gene (Asian pears [Pbr013451], European pears [Pbr039977.1]) that encode sorbitol transporter was also discovered in pear during the large-scale resequencing of pear germplasm (Wu et al., 2018).

Gene duplication is a major mechanism for biologic anagenesis and gene function evolution, providing abundant evolutionary materials for species (Zhu et al., 2021). Gene duplication events can be classified as small-scale (tandem duplication, proximal duplication, dispersed duplication, and transposition duplication) or large-scale (Whole-genome duplication, WGD) (Reams & Roth, 2015). Tandem and proximal duplicates occur much more frequently than WGD, and are believed to maintain a relatively stable state, evolving some plant-specific functions under strong selection pressures (Qiao et al., 2019). Tandem duplication produces a large number of gene copy number and allele variation in a population, and is often important for plants to adapt to rapidly changing environments (Hanada et al., 2008). Different theories related to multiple copy genes in various species, stages and genotypes, including the “avoidance of gene dosage imbalance” model, “duplication, degradation and complementation” model, or “drift compensation” model, all affirm the importance of gene duplication in the origin and development of species and evolution of genomes (Zhang et al., 2019).

The PLT, which includes SOTs, has undergone expansion, particularly through tandem duplication in *Gramineae* crops (Kong et al., 2020, Johnson & Thomas, 2007). Duplication and amplification of SOTs have also been observed in pears, with single-gene duplication being the main mechanism (Qiao et al., 2018). Studies on gene duplication types have shown that WGD, tandem, and proximal duplications play a role in the expansion of peach SOTs, while only dispersed duplication was detected in apple and strawberry (Yu et al., 2019). Despite previous studies on SOTs and their gene family duplications, the origins and evolution of SOTs in plants remain largely mysterious, and the complicated relationships between SOT orthologs and paralogs resulting from abundant duplication require further exploration. A systematic comparison of the syntenic relationships among all SOT genes across multiple plant species can help to shed light on this problem. Synteny information is an essential component of comparative genomics analysis, as it allows for the analysis of both large scale (e.g., genome rearrangement and duplicate events) and small scale (e.g., genome-level base substitution rate and insertions and deletions) molecular evolution events between species, particularly in revealing key events in gene duplication and transposition (Kerstens et al., 2020). Synteny networks (Zhao & Schranz, 2017) are effective tools for organizing and utilizing collinearity data between genomes of numerous sequenced species, enabling the identification of previously undiscovered and complex evolutionary relationships in various gene families. Examples of gene families using this approach include MADS-box (Zhao et al., 2017), APETALA2 (Kerstens et al., 2020b), CBLs and CIPKs (Zhang et al., 2020). Thus, network clustering is a promising method for studying synteny of gene families concurrently in multiple species, and can help to organize and interpret massive pairwise syntenic relationships.

In this study, we analyzed the synteny relationships of all identified PLT genes in 61 angiosperm genomes and SOTs genes in major representative Rosaceae genomes. We combined our finding with phylogenetic analysis to investigate the lineage-specific expansion and syntenic conservation of PLT and SOTs across difference plant lineages. Additionally, by comparing the duplication differences of different SOT clades, we deduced the complex gene duplication pattern of SOTs in Rosaceae, which had not been previously detected. Our study revealed the early origin and evolution of SOT genes from an evo-devo perspective.

## Results

### 1. The PLT gene family in plants exhibits differential expansion across lineages

PLT is the closest evolutionary relative of the SOT gene family, which is considered an important component of PLT. To understand the early origin and evolution of SOT genes, we expanded our investigation from SOT to PLT. We constructed a comprehensive dataset of proteins across 61 major representative plant lineages to investigate the distribution of PLT in angiosperms. Initially, we used the conserved domain (Sugar_tr, PF00083) of PLT to construct the hidden Markov model (HMM) and searched against the dataset. Based on the mapping results and taxonomy annotation of the obtained sequences, we found that Sugar_tr is not a PLT-specific domain (Supplemental Figure 1). Although we retrieved the PLT sequence, other non-PLT sequences also appeared. The majority of candidate sequences belonged to the MST gene family, which consists of seven gene families that include PLT. The remaining sequences belonged to other non-MST gene family from the MFS superfamily, such as phosphate transporters and carnitine/organic cation transporters (OCTs). Therefore, we used sequence alignment to identify PLT specificity determining regions and built a PLT-targeting HMM (Supplemental Figure 2). With this model, we could extract PLT sequences intact from the primary screening dataset of PF00083.

A total of 704 PLT amino acid sequences were identified across 61 plant genomes covering diverse angiosperm groups, and a phylogenetic tree was constructed (Figure 1A). Phylogenetic analyses revealed that PLT genes can be divided into three solid clades with high bootstrap support: the Class I, Class II, and Class III. Within Class III, four distinct subclades can be distinguished: Class III-A, Class III-B, Class III-C, and Class III-D. The PLT gene family in plants exhibited differential group expansion across lineages (Figure 1).

**Figure 1.**
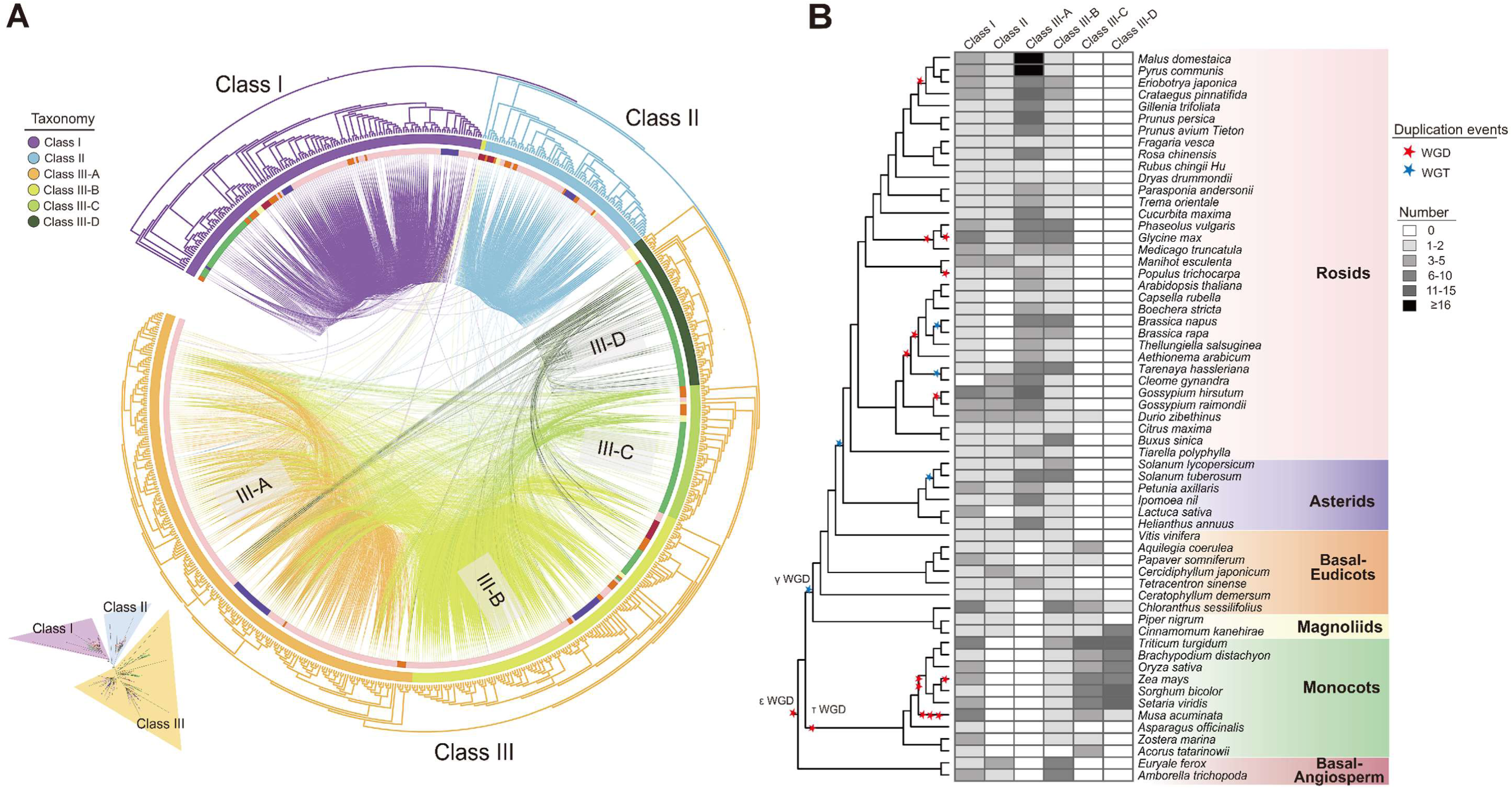
Phylogenetic tree and distribution of the PLT family in Angiosperms. **(A)** Maximum-likelihood gene tree for the PLT gene family and syntenic relationships between the genes. Terminal branch represent three broad categories: Class I, Class II, and Class III. The first inner circle represents six subgroups: Class I, Class II, Class III-A, Class III-B, Class III-C, and Class III-D. The second inner circle represents genes belonging to Rosids (light pink), Asterids (blue), monocots (green), Magnoliids (yellow), basal-eudicots (orange), and basal-angiosperm (red). Each connecting line located inside the inverted circular gene tree indicate strong conservation of synteny between gene pairs within subclades. The connecting lines are colored according to the corresponding communities. Bottom left: unrooted view of the major clades of the complete PLT tree. **(B)** The copy numbers of Class I, Class II, and Class III are shown for each plant species. The degree of enrichment is represented by the depth of the gray-colored cells. Red and blue pentagrams on the tree indicate known whole-genome duplication (WGD) and whole-genome triplication (WGT) events, respectively, the phylogenetic tree on the left side.

Class I was a basal clade that included 145 sequences from all plant species in this study. The eudicot members in Class I could be devided into two subclades, and a strong duplication signal resulting from WGDs were found between the two clades (Supplemental Figure 3). Class II consisted of 78 sequences from 43 angiosperm species. The members of Class II were all from eudicots (except Brassicaceae) and basal angiosperms (Figure 1B). In other words, Class II was an angiosperm branch that completely lacked monocot sequences. In the phylogenetic tree, a large number of tandem pairs were identified in many orders of Super-Rosids, such as Rosales, Malpighiales, Malvales, and Sapindales (Supplemental Figure 4). However, no tandem duplication pair was detected in basal eudicots like *Vitis vinifera* or *Aquilegia coerulea*. The same tandem pairs did not exit in the Super-Asterids, and only a single gene was present in each species. We inferred that Class II underwent specific evolution and may have been lost in the ancestor that gave rise to the monocot lineage. The retained eudicot members experienced duplication and loss, eventually evolving into a lineage-specific branch. To date, few studies have reported on the function of PLT, and the underlying evolutionary and functional effects of species-specific gene deletion in Brassicaceae and monocots await further investigation.

The Class III is composed of 481 angiosperm sequences, with 216, 136, 59, and 60 genes belonging to Class III-A, Class III-B, Class III-C, and Class III-D, respectively (Figure 1A). Class III-A is specific to eudicots, with sequence from *Vitis vinifera*, *Cercidiphyllum japonicum*, and *Tetracentron sinense* (basal-eudicots) also detected (Figure 1B). The number of sequences in Class III-A is significantly higher than that of the other groups, accounting for nearly one-third of the total number of PLT sequences. This suggests that Class III-A group flourished well during plant evolution. Interesting, a large number of homologous genes were observed in Rosaceae and especially in the Amygdaloideae in Class III-A. The clade formed by Rosaceae species could be further divided into two subclades with strong support (Bootstrap = 91%) (Figure 3A).

Class III-B is a complete species composition branch, with sequence from almost all angiosperm species of this study detected. Two clades, called Class III-C and Class III-D, with a strong preference for monocots, were isolated from Class III. The sequences of monocots were enriched to a greater degree than the other clades (Figure 1B).

### 2. Conservation and Lineage-Specific Synteny Relationships in PLT

The similarity of gene arrangement on chromosomes in different genomes, known as synteny, can be used to compare the rules (conservation or lineage-specific) of gene arrangement at the level of genome structure and infer the shared ancestry of genes (Gabaldon & Koonin, 2013, Zhao & Schranz, 2019). To understand the evolution history of PLT, we constructed a phylogenomic synteny database of 61 species and screened for PLT-targeting synteny networks. We conducted a phylogenomic analysis of PLT by combining phylogenetic tree and synteny network. Of the 704 PLT sequences, 524 PLTs could be linked with at least one other homologous seqeuence through synteny. The internal homologous pairs of each major clade showed a highly syntenic connection signal. This syntenic signal supports the reliability of classifications of the PLT family in angiosperms from the phylegenetic analysis. Three major clusters were identified on the synteny network (Figure 2A).

**Figure 2.**
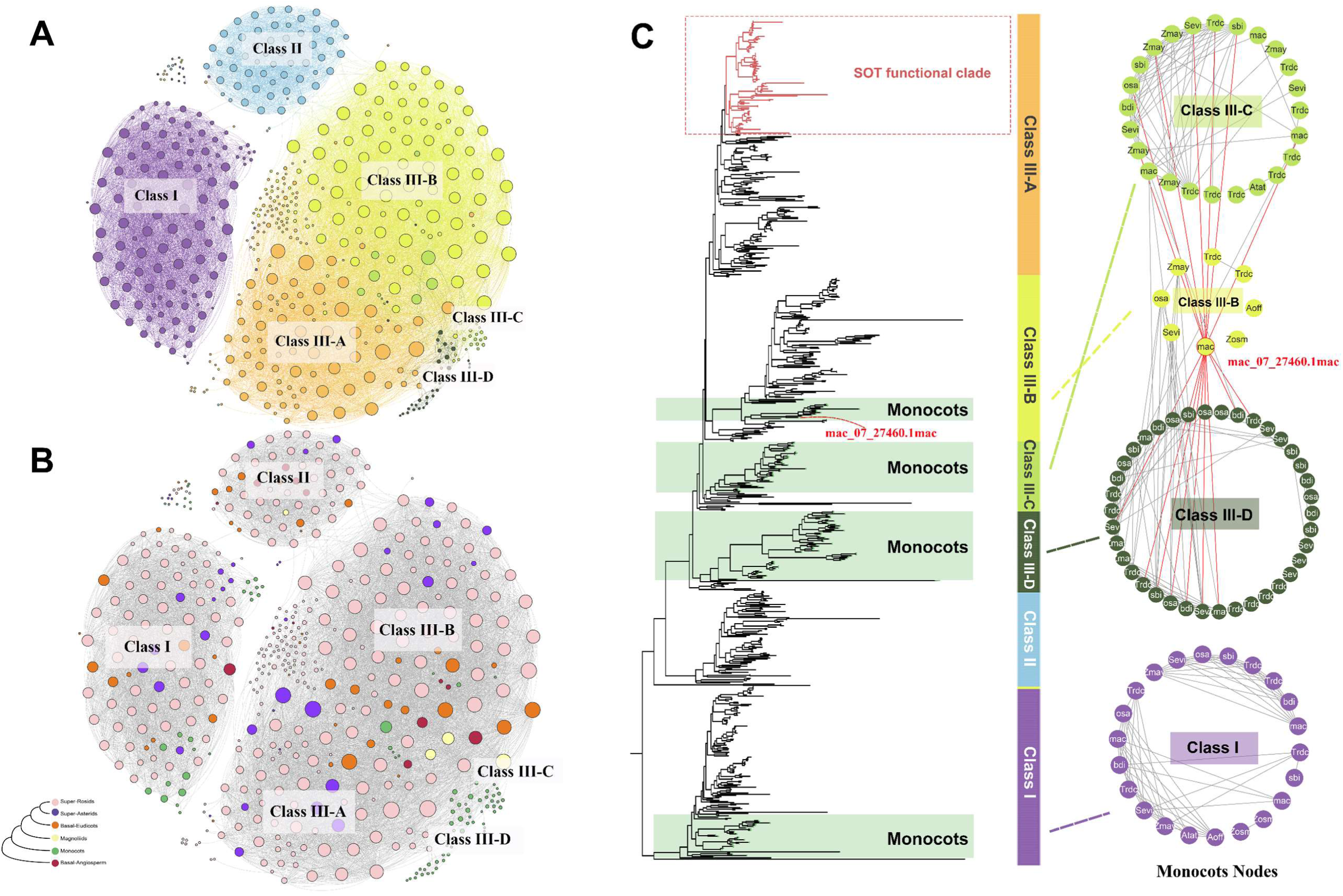
Syntenic relationships of the PLT family in angiosperms and detailed network representation for a selected set of important monocots synteny clusters. **(A)** Synteny network of the PLT gene family using all the detected syntenic relationships in the synteny network database. Communities were generated based on the clique percolation method at k= 3. The size of each node corresponds to the number of edges it has (node degree). Nodes are colored based on the PLT subfamilies, which is consistent with Figure 1A. **(B)** Nodes are colored based on the taxa they belong to. Communities are indicated by the corresponding subfamilies. The rest of the visualization is the same as in A. **(C)** Monocots nodes were extracted from the entire phylogenomic synteny network and formed a sub-network. The nodes were laid out and colored according to their corresponding categories. The colors used are consistent with (A). The left panel shows the distribution of monocot sequences on the PLT tree. The clades representing functional SOT genes are highlighted in dull red.

The sequences of Class I formed a separate cluster, with almost no syntenic signal between Class I and other clade. Given its limited synteny communities and ubiquitous feature, we speculated that this clade has a conserved and essential basic function.

The synteny network revealed that two gene clusters were enriched by Class II and Class III, respectively. The small cluster was composed of Class II genes, while the large one was enriched by Class III. The Class II genes showed weak synteny with Class III (Figure 2A and B). Based on the similarity of the branches on the phylogenetic tree, we infererred that Class II is more closely related to Class III as they belong to sister clades, whereas the clade of Class I is relatively distant from them. Furthermore, when combined with the collinearity of *Amborella trichopoda* and *Euryale ferox* (basal-angiosperm), it can be inferred that PLT gene in the common ancestor of Class I, Class II, and Class III may have undergone an ancient whole-genome-duplication event in angiosperms, leading to the emergence of three different gene clusters (Supplemental Figure 5A).

The strong synteny network consists of Class III-A, Class III-B, Class III-C, and Class III-D (Figure 2A and B), and the node number of this network accounts for more than half of the entire network. Based on the mapping and taxonomy annotation of each node, some interesting phenomena were observed in this strong synteny cluster. The nodes at the center of the cluster were composed of basal-angiosperm, magnoliids, and basal-eudicots sequences. On both sides of the network were a large number of nodes from super-Rosids and super-Asterids, which belong to the Class III-A and Class III-B branches. The composition and structure of species suggest that Class III-A genes are closely related to Class III-B genes.

Further investigation into the synteny relationships of the genes of basal-eudicots (Supplemental Figure 5B) revealed that the homologs from Class III-A and Class III-B (vvichr3G00005650, vvichr4G00009460) demonstrated collinearity. Combined with phylogenetic tree, it is suggested that genes of Class III-A and Class III-B are derived from an ancient common ancestor, and then Class III-A expanded by gamma hexaploidization (referred to as γ triplication) of eudicots.

Furthermore, except for eight nodes in Class III-B, the remaining monocot species in Class III form two separate small synteny clusters attached to the edge of the strong network (Figure 2A and B). We performed association analysis for the nodes using phylogenetic tree to confirm that the nodes belong to Class III-C and Class III-D repectively. To disentangle the associations of monocots in Class III, monocot nodes were extracted from the whole phylogenomic synteny network and formed the sub-networks (Figure 2C). The related nodes fall into two separated subgroups. The monocots homologs of Class III-B, Class III-C, and Class III-D were clustered, with Class I proteins of monocots independently forming a single subgroup in the subnetwork (Figure 2C). These synteny patterns suggest that Class I and Class III may have different origins. One node (mac_07_27460.1mac) belonging to Clssss III-B was detected and showed synteny relationships with Class III-C and Class III-D simultaneously. We suggest that the monocots in Class III formed three groups due to the ancient whole genome dupliation (WGD) shared by all monocot species, followed by intraspecific duplication and differentiation.

### 3. SOTs in Rosaceae evolved from a pair of PLT Class III-A tandem duplicated genes

Next, we used a phylogenomic approach to explore the origin and evolutionary history of SOTs. We used the protein sequences of reported SOTs (Supplemental Table 3) as query sequences to retrieve the homologous genes from the available PLT protein dataset. We found that all reported SOT sequences are located in the Class III-A branch. However, species outside of Rosaceae do not primarily produce sorbitol as a photosynthetic product and only contain trace amounts of sorbitol or are unable to synthesize it. Therefore, the physiological functions of Class III-A genes in non-Rosaceae plants are likely different from those in Rosaceae, where the transport function of these PLTs is more extensive, even though PLTs in non-Rosaceae may not transport sorbitol in the body. For example, cloning AtPLT5 from *Arabidopsis* and expressing it in *Xenopus oocytes* proved that AtPLT5 transports a wide range of hexoses, pentoses, and sugar alcohols including sorbitol, but it may not translocate or accumulate sugar alcohols such as sorbitol in *Arabidopsis* in vivo (Reinders et al., 2005, Klepek et al., 2010). Further studies revealed that the Class III-A genes in Rosaceae, defined as “SOT” in previous studies, except for one gene from *Dryas drummondii* (Drydrscaffold178G00001580), are different from those in non-Rosaceae. As the primary carrier of sorbitol translocation from source to sink, SOTs in Rosaceae have become more specialized in sorbitol transporting.

As the basal species of Rosaceae, two tandem copies (Drydrscaffold178G00001580, Drydrscaffold178G00001630) were found in the Class III-A branch (Figure 3A). BlastP analysis revealed that one copy (Drydrscaffold178G00001630) more closely resembled SOT homologs, while the other copy (Drydrscaffold178G00001580) was more similar to Class III-A genes. Based on the phylogenetic tree, Drydrscaffold178G00001630 clustered with other sequences from Rosaceae (Figure 3), while Drydrscaffold178G00001580 was isolated and grouped into a separate branch, indicating a relatively distant evolutionary relationship between the two genes.

**Figure 3.**
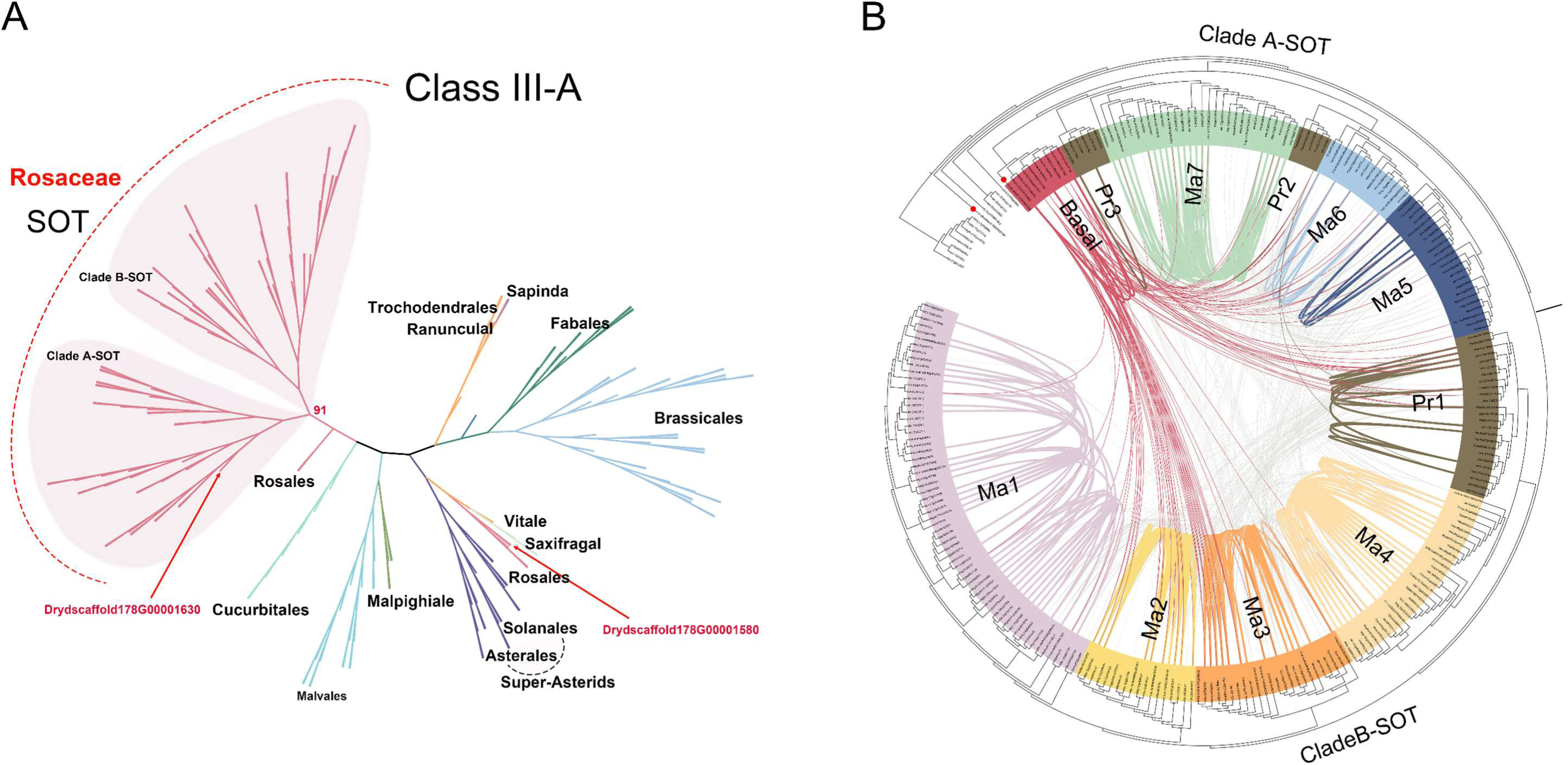
Maximum-likelihood gene tree and syntenic relationships for the SOT gene family. **(A)** Detailed species distribution for the Class III-A clade, which contains the SOT clade. **(B)** Colored lines (except red) inside the inverted circular gene tree indicate strong conservation of synteny between gene pairs within subclades, while gray lines indicate syntenic connections between subclades. Red lines indicate strong conservation of synteny between gene pairs within the Basal subclade, as well as between Basal and other subclades. Brown lines indicate strong conservation of synteny within Pr subclade. The Drydrscaffold178G00001580 (left red dot), Drydrscaffold178G00001630 (right red dot) are highlighted separately.

We used synteny network to verify the phylogenetic relationship between Class III-A and SOT. The SOTs showed a strong syntenic signal with other Class III-A genes in non-Rosaceae, and we also found that Drydrscaffold178G00001580 could form syntenic pairs with SOT genes in Rosaceae. These results suggest that SOTs in Rosaceae evolved from a pair of Class III-A tandem duplicating genes and underwent a complex history of functional evolution.

### 4. Identification and phylogenetic reconstruction of the SOT gene family in Rosaceae

As discussed earlier in the previous section, the number of PLTs in Rosaceae is far greater than in other species. After an initial investigation, we found that a substantial number of SOT gene duplications is the main reason for this phenomenon. To further investigate the process and nature of gene expansion that occurred in the SOTs, we identified SOT protein sequences from representative Rosaceae genomes, including Amygdaloideae (*Malus domestaica* [apple], *Pyrus communis* [pear], *Eriobotrya japonica* [loquat], *Crataegus pinnatifida* [hawthorn], *Gillenia trifoliata* [Bowman’s root], *Prunus persica* [peach], *Prunus mume* [plum], *Prunus armeniaca* [apricot], *Prunus avium* [sweet cherry]), Rosoidaea (*Fragaria vesca* [strawberry], *Rosa chinensis* [rose], *Rubus occidentalis* [raspberry]) and *Dryas drummondii* (Dryadoideae). We then performed further phylogenetic analysis uisng SOTs protein sequences from the above Rosaceae species (Figure 3B). We found that the SOTs cluster into two major groups with strong support based on sequence similarity and phylogenetic analysis. One clade is a basal clade that incorporates proteins from Amygdaloideae, Rosoidaea, and Dryadoideae, termed “Clade A-SOT”. The other clade is an Amygdaloideae-specific clade with no proteins from the Rosoidaea and Dryadoideae lineage, termed “Clade B-SOT”. The branch in Figure 3B corresponding to the branch in Figure 3A.

The two SOT clades identified in Rosaceae could be further divided into 11 groups based on species and bootstrap support, designated as “Basal”, “Pr1” to “Pr3”, and “Ma1” to “Ma7”. The basal clade includes SOT genes from both Dryadoideae and Rosoideae species, while the remaining groups were derived from Amygdaloideae. “Pr1” to “Pr3” corresponding to the SOT sequences from Prunus species, while “Ma1” to “Ma7” contain the sequences from Maleae and Gillenieae species.

Gene duplication is a major mechanism of biological anagenesis and gene function evolution. SOTs in Rosaceae exhibit a high incidence of lineage-specific expansions. However, no large-scale SOT gene expansion patterns were observed in Dryadoideae and Rosoideae, where only one or two SOT gene was identified. In contrast, the SOT family has an extensive duplication history in Amygdaloideae species, with Maleae being the center of gravity for duplication. In particular, five copies were found in rose, implying a gene duplication event occurred after speciation.

### 5. Synteny relations across Amygdaloideae for overlooked but interesting SOT gene

To understand how the SOT gene family expanded in Amygdaloideae, we analyzed the expansion history of the gene family based on both the phylogenetic trees and chromosome localization in four representative species. We also examined potential SOT gene duplications and losses in Amygdaloideae by analyzing the degree of local synteny at the SOT locus.

In apple, a total of 19 SOT genes were mapped to Chromosome 5 and Chromosome 10, except for two (Figure 4A). A similar pattern was observed in loquat, where two chromosomes harbored all the SOT genes except for one gene mapped to Chromosome 17 (Figure 4B). Thus, the distribution of SOTs on chromosomes was consistent in Maleae species, with most of genes concentrated on just two chromosomes. SOT genes on the same chromosome tended to be tandemly clustered. Notably, combining the phylogenetic tree with chromosomal localization, we found that most paired genes, which belong to the same subfamily, were distributed on different chromosomes. The sequence identity between member pairs on different chromosomes was higher than that within the tandem cluster on the same chromosome. We performed microsynteny analysis and found that in apple, the locus containing a SOT homolog on Chromosome 5 showed high-level synteny to the paired gene locus of Chromosome 10, which belonged to the same SOT families (Figure 6A). This synteny pattern was also found in loquat. These findings suggested that paired genes from the same subfamily but located on different chromosomes have higher homology and are more closely related than tandem genes from different subfamilies on the same chromosome. Based on these results, we inferred that the ancestor genes of each SOT subfamily in Maleae underwent divergence events before WGD events.

**Figure 4.**
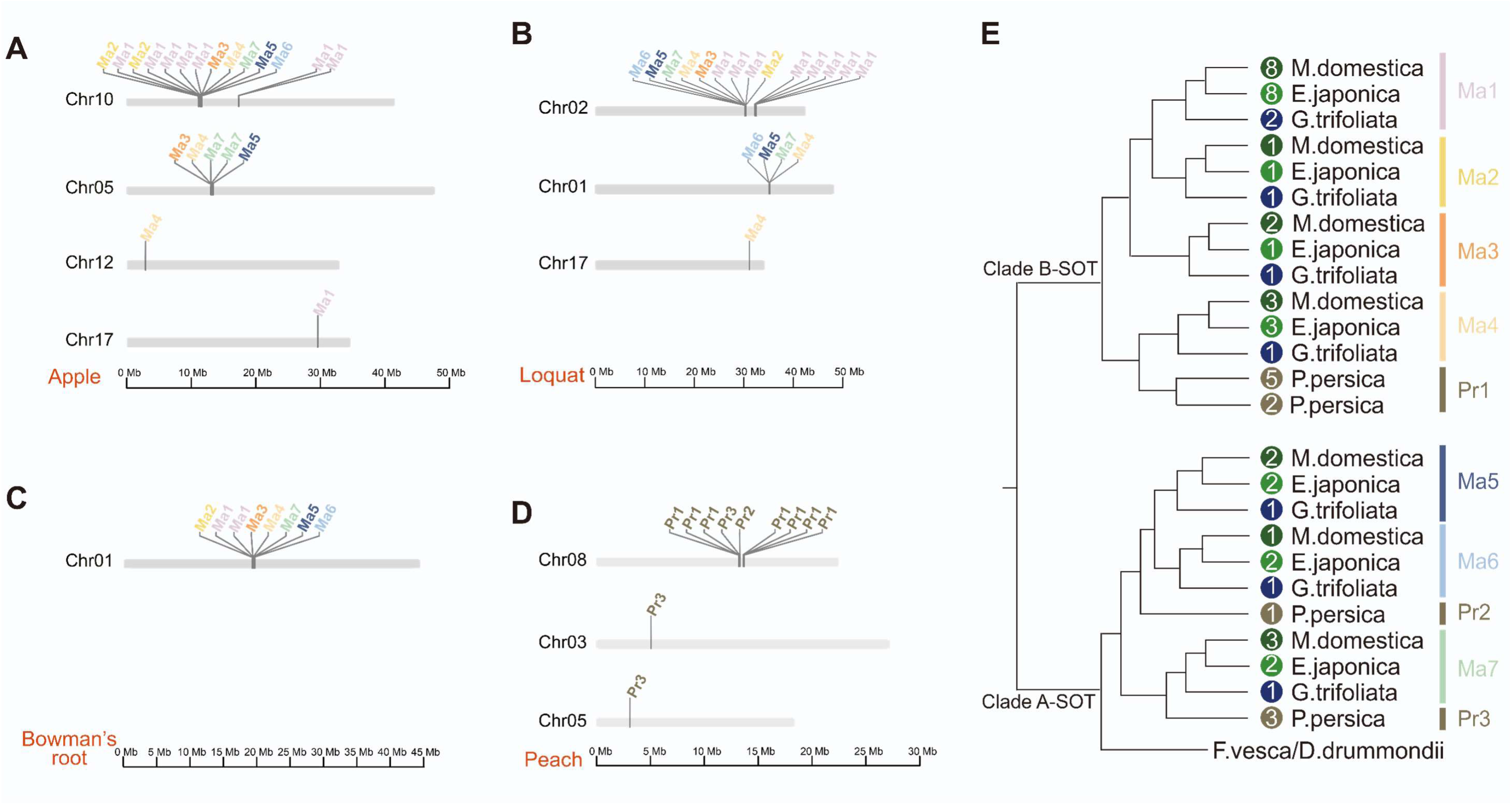
Chromosomal distributions of the identified SOT genes. Chromosome numbers are shown on the left of each bar. The scale at the bottom indicates chromosome sizes. Panels (**A-D**) show the distributions of SOT genes in apple, loquat, bowman’s root, and peach, respectively. Each colorful label represents a gene member from different subfamilies, and the colors were colored according to the subfamilies identified in Figure 3(B). **(E)** Copy number changes of SOT in four Amygdaloideae species, with circle numbers representing the numbers of SOT genes in each species.

Gillenieae is regarded as the sister group of Maleae, as its genome has not been affected by most recent WGD shared by Maleae. The representative species in Gillenieae is bowman’s root. In the phylogenetic trees, unlike Maleae species with multiple copies of orthologs in the same subfamily, bowman’s root had only one sequence in each subfamily, except the SOT1 subfamily (Figure 4C). All SOT genes were located in a single cluster on Chromosome 1. Interestingly, SOT homologs in bowman’s root and Maleae species are highly syntenic within their subfamilies and show a one-to-many species sequence distribution pattern between bowman’s root and Maleae (Figure 4E). These phenomena suggest that the subfamily divergence of SOTs can be traced back to the time before the divergence of Maleae and Gillenieae.

Peach and bowman’s root share a similar pattern in which nearly all SOT genes were located on the same chromosome and distributed in a tandem manner (Figure 4D). The phylogenetic tree and chromosome location showed that all sequence in peach were expanded by tandem duplications, except the ancestor of a cluster including four tandemly arrayed SOTs, which underwent a unique segmental duplication event.

In total, SOT homolog was identified in the syntenic regions of apple, loquat, bowman’s root, and peach. Amygdaloideae showed a high degree of genomic synteny. The locus of SOT homologous genes from the same subfamily in the four species showed high local synteny among each other.

### 6. Noteworthy Lineage-specific gene duplications and transpositions of SOT in Maleae

The replication mode of SOT1 was found to be more complex than other subfamilies, with the SOT1 gene cluster being the most numerous. While only two tandem SOT1 genes were detected in *G. trifoliata*, Maleae species showed a large number of SOT1, indicating that SOT1 underwent additional lineage-specific tandem expansion in Maleae (Figure 4). Moreover, two SOT1 tandem genes (Mgde10G1106100 and Mgde10g1106200) located on Chromosome 10 in apple were distant from most of the SOT tandem clusters. Although the four genes up- and downstream of SOT1 were highly collinear in all four Amygdaloideae species, these two genes could not be detected as collinear orthologous genes via local synteny analysis in loquat, bowman’s root, and peach (Figure 5D). In a SOT-targeting synteny network of 19 Rosaceae species, the two SOT1 genes that were speculated to be obtained by fragment duplication and located in a separate small network, which was formed only by SOT1 genes from various apple cultivars (Figure 5A and 5B). Based on the collinearity of chromosomal regions containing the two genes and the formation of this specific synteny network, we interpreted that these genes might be the products of intraspecific duplication and transposition events.

**Figure 5.**
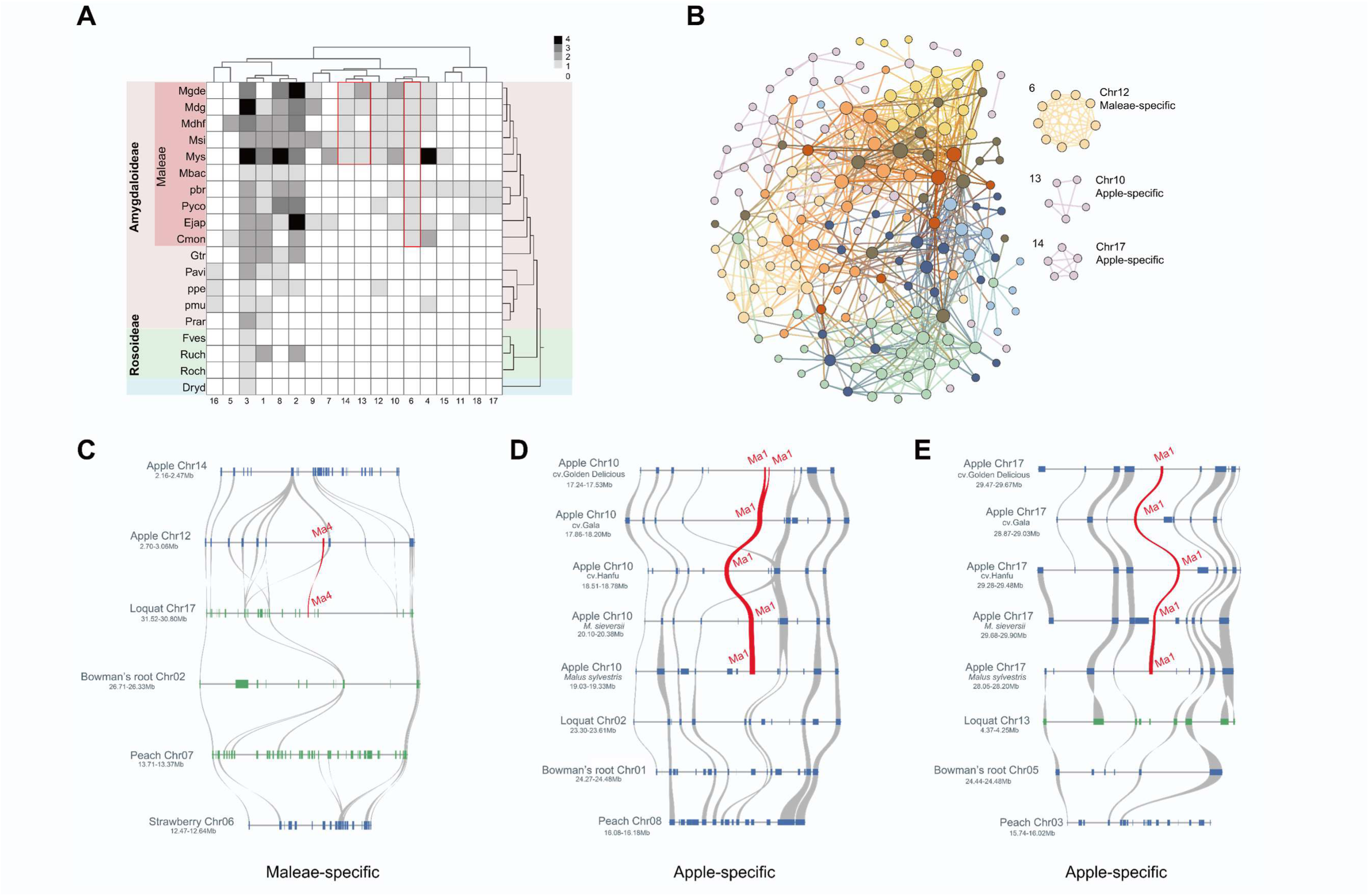
Phylogenetic profiling and synteny relationships of the SOT homologous genes in Rosaceae. **(A)** Phylogenomic profile of SOT syntelogs (syntenic homologous genes) across Rosaceae genomes. Species names are shown on the left side with their corresponding abbreviations as listed in Supplementary Table 2. The corresponding phylogenetic tree of species is shown on the right side. The cluster ID is indicated at the bottom of the figure. The columns show syntelog clusters, and cell shading indicates the number of nodes (genes grouping in that cluster) per species. **(B)** The synteny network of the SOT gene family in Rosaceae species. Clusters from (A) can be divided into large-conserved clusters and lineage-specific clusters, including Maleae-specific (Cluster 6) and Apple-specific (Cluster 13–14). The size of each node corresponds to the number of edges it has (node degree). The nodes are colored according to the discovered SOT subfamilies in Figure 3(B). **(C)** Synteny relationships of the Maleae-specific SOT locus region in apple and homologous segments in other species from Amygdaloideae. Grey curves connected the identified syntenic genes, while the annotated genes are represented by rectangles. Genes located on the forward strand are shown in blue and genes located on the reverse strand are showed in green. SOT homologs are marked in red and connected by red lines. **(D-E)** Microsynteny relationships of the apple-specific SOT locus region in apple and homologous segments in other species from Amygdaloideae.

Using similar approaches, we found that the SOT1 gene in apple, located on Chromosome 17, was most likely the product of intraspecific single-copy and experienced relocation after the WGD (Figure 5E).

Interestingly, in addition to the SOT4 duplication gene shared by all Amygdaloideae species, a unique random duplication of the SOT4 gene was found in Maleae plants. A small lineage-specific synteny network consisting of only SOT4 single-copy from Maleae species demonstrated the existence of this specific gene (Figure 5A and B). Further microsynteny analysis showed that the SOT4 gene, located on Chromosome 12 in apple, had a strong syntenic relationship with SOT4 gene on Chromosome 17 in loquat (Figure 5C). No gene corresponding to this specific copy gene was found in bowman’s root and peach, implying that this gene may have arisen from a unique random duplication in the ancestor of Maleae and occurred after the recent independent WGD events.

### 7. Expression profiles of SOT genes in Rosaceae

To further study the functions of SOT genes in Rosaceae, we analyzed their tissue-specific expression in apple, bowman’s root, and peach (Figure 6B). Differential expression analysis revealed the transcriptional differences among duplicated genes, with the Clade A-SOT genes showing higher expression levels than the Clade B-SOT genes. Different members of SOT gene families exhibited tissue-specific expression patterns. The Ma1 genes were completely inactive, indicating that they have lost function. The Ma2 genes were expressed specifically or preferentially in flowers, suggesting that they may play crucial roles in flower development. The Ma4 genes show higher expression in leaves compared to other tissues, while the Ma6 genes were ubiquitously and highly expressed in all tissues. Therefore, it is inferred that Ma6 may play a critical role in sorbitol transport and sugar metabolism. Although some homologous genes exhibited similar expression patterns, there are considerable differences in the degree of expression. These results suggest that SOT members may have functional redundancy.

**Figure 6.**
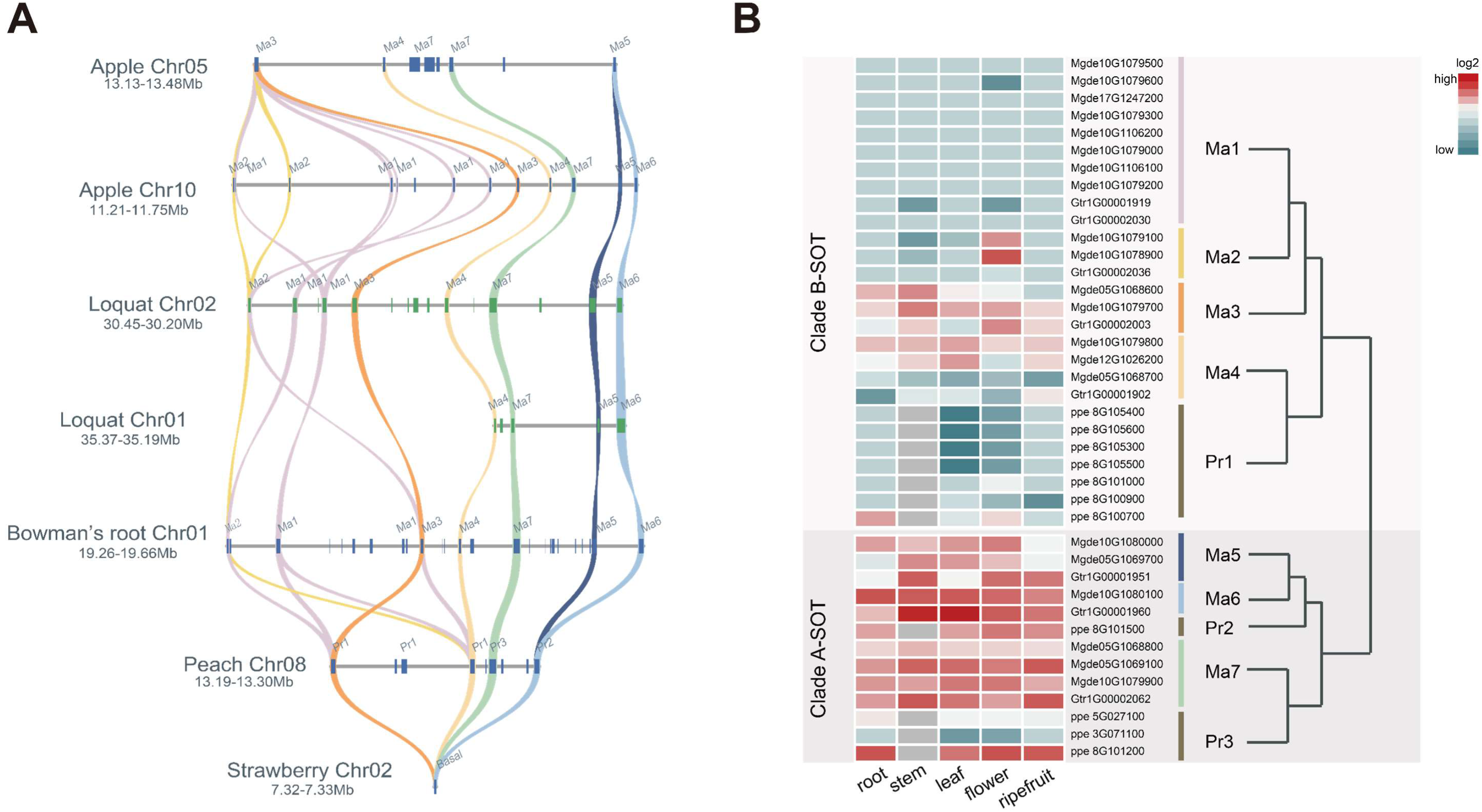
Microsynteny relationships of SOT homologous segments in Rosaceae species and tissue-specific expression patterns of SOTs. **(A)** Syntenic relationships at the SOT loci were showed in Rosaceae species, including apple, loquat, bowman’s root, peach, and strawberry. Curves connect the identified syntenic genes, with labels on the rectangle indicating SOT genes. SOT homologs belonging to same subfamilies are marked in same color and connected by same color lines, as determined by the subfamilies discovered in Figure 3. **(B)** The order of sequences matched the SOTs in the phylogenetic tree shown to the right of the expression profiles. Gray rectangles indicated missing data for peach. Species abbreviations are as follows: Mgde, *Malus domestica*; Gtr, *Gillenia trifoliata*; ppe, *Prunus persica*.

### 8. The evolutionary model of SOTs

Based on the results presented above, we proposed an evolutionary history of the SOT gene family in Rosaceae (Figure 7). The tandem gene arrangement pattern appears to have arisen from a common ancestor of the Amygdaloideae, which experienced two consecutive tandem duplications of SOT genes. The first duplication event resulted in the formation of the Clade A-SOT and Clade B-SOT ancestral genes, while the second duplication event allowed the separation of Ma7 and Ma5/6 in Clade A-SOT as well as Ma3 and Ma4 in Clade B-SOT. Subsequently, a series of tandem duplications occurred after the divergence of Maleae and *G. trifoliata* from the Amygdaloideae, resulting in the formation of the current Ma1-7 tandem gene arrangement. A recent WGD event occurred prior to the diversification of extant Malea, allowing for the emergence of paired genes from the same subfamily but located on two different chromosomes. Finally, intraspecific single-gene losses and lineage-specific transpositions were found.

**Figure 7.**
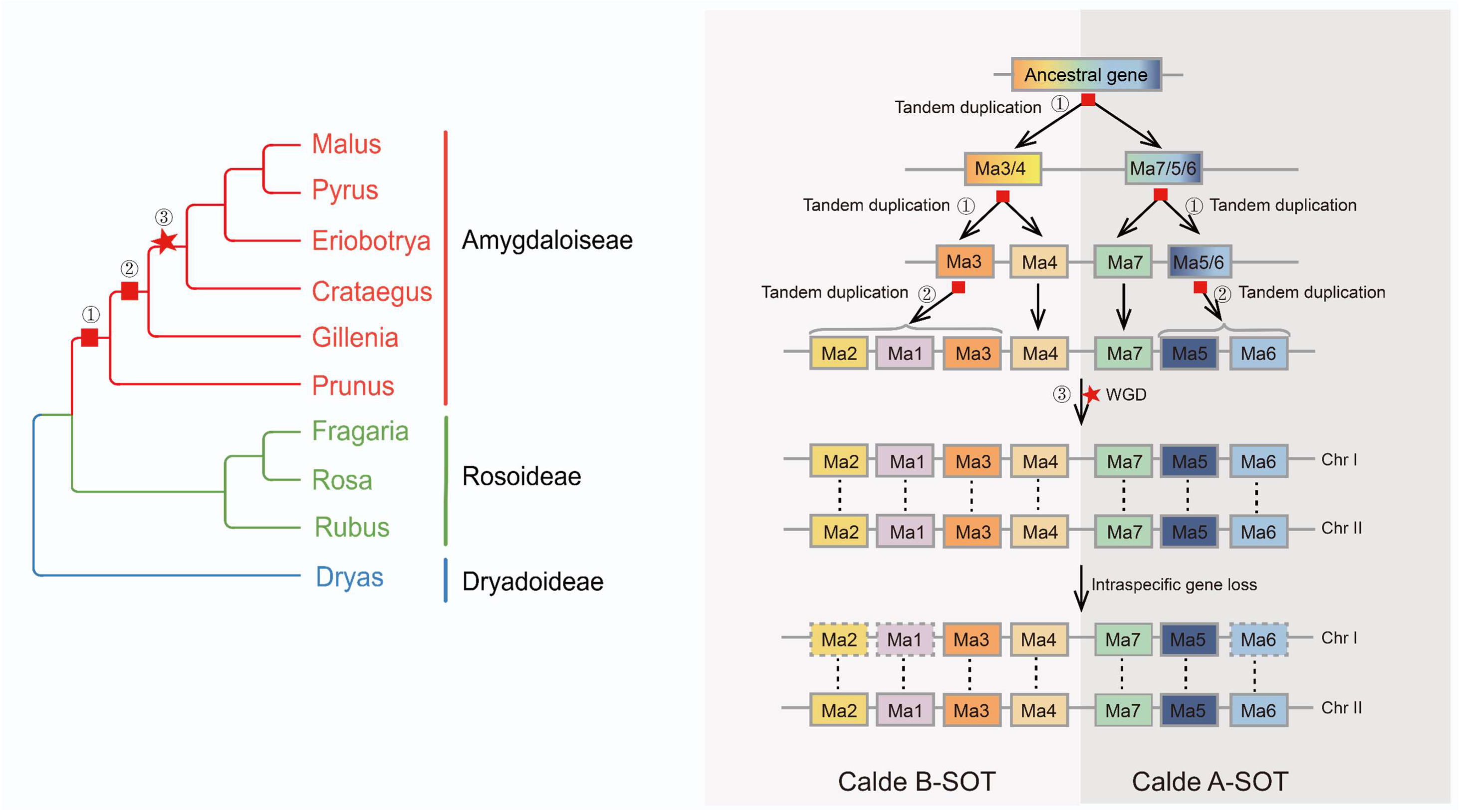
The evolution scenario of the SOT gene family. Rectangles of the same color with label represents the same subfamilies, and the colors were colored according to the subfamilies identified in Figure 3(B). Red rectangles and pentagram indicate tandem duplication and whole-genome duplication (WGD) event, respectively, the corresponding phylogenetic tree of species is shown on the left side.

## Discussion

Sorbitol have been recognized as an important photosynthesis-derived carbohydrates that plays a role in growth, development, and resistance of Rosaceae (Gao et al., 2003, Aprea et al., 2017). Sorbitol is mainly transported and mediated by SOTs, which are one of the important components of the PLT family (Fan et al., 2009). The origin and evolutionary history of SOT gene family duplication in angiosperms remain unclear. In this study, we investigate the synteny relationship of all the detected Polyol/Monosaccharide Transporter (PLT) genes in 61 angiosperm genomes and SOT genes in major representative Rosaceae genomes. Through a combination of phylogenetic analysis and comparative genomics, we aim to elucidate the lineage-specific expansion and syntenic conservation of PLTs and SOTs across different plant lineages and to propose a gene duplication model of SOTs that can provide insights into the genomic origins, duplication, and divergence of SOT gene family members.

### 1. The Ancient Origin and Expansion of the PLT Family in Plants

The PLT gene family, to which SOT belongs, has been demonstrated to transport polyols and monosaccharides (Klepek et al., 2010). To investigate the early origin and evolution of SOT genes, we extended our focus from SOT to the PLT gene family. In our exploration of the PLT gene family in angiosperms, we used phylogenetic analysis that combined phylogenetic reconstruction and synteny networks. Based on this analysis, the PLT gene family could be divided into three broad categories (Class I, Class II, and Class III) and six subgroups (Class I, Class II, Class III-A, Class III-B, Class III-C, and Class III-D).

Class I genes were found to span a broad range of angiosperm lineages. We further exploited the PLT-targeting Hidden Markov model we created to perform a genome-wide screen for PLT in species outside angiosperms, including *Ginkgo biloba* (gymnosperms), *Salvinia cucullate* (ferns), *Azolla filiculoides* (ferns), *Marchantia polymorpha* (Mosses), *Physcomitrella patens* (mosses), and *Chlamydomonas reinhardtii* (chloraphyta). All lineages except for *Chlamydomonas reinhardtii* contained PLT. The close clustering of the selected sequences of these species with Class I suggests a closer phylogenetic relationship (Supplemental Figure 6). Class I genes were found in broad-scale lineages spanning from early-diverging mosses to angiosperms, suggesting an ancient origin and implying that this clade has a conserved and essential basic function.

The PLT family is an ancient and widely distributed protein transporter family, with origins dating back to the fern lineage (Johnson et al., 2006). Our analysis confirmed the absence of PLT in charophyta (*Chlamydomonas reinhardtii*), while the identification of a single PLT gene in *Physcomitrella patens* and *Marchantia polymorpha* suggests that the origin of PLT genes may trace back to mosses. This observation indicates an earlier origin of the PLT genes than previously reported, which broadens the scope for future research on PLT.

Our analysis of the Class II subfamily genes revealed a striking anomaly – these genes were completely absent in Brassicaceae and monocots, indicating lineage-specific gene loss. While we are unable to identify a specific trait of Brassicaceae and monocots that correlate with this phenomenon due to the limited studies on the function of PLT family members, this observation aligns with the birth-death model of gene family evolution, in which gene duplication creates new genes that may be maintained or lost over time (Panchy et al., 2016). Future functional analysis is needed to confirm this hypothesis.

Class III-A is the most extensively evolved branch in the PLT family, with a substantial number of homologous genes present in Rosaceae. Unlike most terrestrial plant, sorbitol is the principal photosynthate and soluble storage material in Rosaceae (Wang et al., 2016, Li et al., 2012). The high demand for large quantities of sorbitol transport may have driven the continuous evolution of PLT protein sequences in Rosaceae. This allowed for the transport of sorbitol characteristics of certain PLT genes to become increasingly prominent, ultimately leading to the development of SOT. Sequence alignment and evolutionary analysis classified the Rosaceae homologous genes clustered in Class III-A as unique to Rosaceae and designated as SOT. These sequences displayed high collinearity with Class III-A sequences of non-Rosaceae plants.

Of note, we identified a tandem gene duplication *D. drummondii,* with one copy (Drydrscaffold178G00001630) belonging to SOT, and another copy (Drydrscaffold178G00001580), which resembles PLT homologs, only appears in the Dryadoideae lineage with no protein from Rosoidaea and Amygdaloideae. The tandem gene duplication event suggests that SOTs in Rosaceae originated from a pair of Class III-A tandem duplicating genes. Following this event, the homologs underwent functional differentiation via gene loss, sub-functionalization or neofunctionalization (Eirín-López et al., 2012). Ultimately, the PLT and SOT genes in Rosaceae diverged.

### 2. Lineage-specific duplication and rapid expansion of SOT

We screened SOT sequences in 19 species genomes, which confirmed all previously identified SOTs in pear and strawberry and identified 11 SOT sequences in peach, nine of which were described before (Gu et al., 2021b, Yu et al., 2019). However, we observed disagreements between our results for apple (Golden delicious) sequence screening and previous results. While previous research showed that the 17 SOT genes of apple were distributed on five chromosomes with enrichment on Chr12 (Yu et al., 2019), our research showed uneven distribution of SOTs with 21 SOTs located on four chromosomes, most of which were on Chromosomes 10 and 5. We attribute the differences to variations in screening approaches and genome annotation versions. To verify the reliability of our results, we tested more apple genomes. The number and chromosome distribution of SOTs were similar in the three cultivated apple species (Golden Delicious, Hanfu, and Gala) and two wild species (*Malus sieversii* and *Malus sylvestris*), after ignoring the segments that have not been assembled onto a chromosome (Supplemental Table 5). These results demonstrate that wild and cultivated species maintained relatively consistent number of SOTs, which proved the accuracy of our identification method.

Gene families occasionally undergone small-scale lineage-specific duplications (Zhang et al., 2020), and several reports have highlighted frequent gene duplications in the SOT gene family (Qiao et al., 2018). Our results are consistent with previous studies, as we found clear evidence that a lineage-specific duplication in Amygdaloideae is the main reason for the expansion of SOTs in the Rosaceae plants.

Regarding gene duplication pattern in Maleae, our study revealed that it is the most drastic and complex compared to other Rosaceae species. Previous studies have shown that the expansion of SOT in pear was dependent on three types of duplication, including dispersed, proximal, and tandem. In contrast, for apple SOT genes, only a discrete replication pattern was found (Yu et al., 2019). However, our study suggests that tandem gene duplication and WGD were the major contributors to the expansion of SOT gene family in Malea. Furthermore, our phylogenetic and synteny analyses showed lineage-specific transpositions, indicating that the gene duplication-transposition could also play a role in the expansion of the SOT gene family in Maleae. The exact cause of the transposition events remain unclear, but it is possible that inter- and intra-chromosomal translocation carrying loci, or the involvement of transposable elements may be contributing factors (Schmitz et al., 2022, Jia et al., 2021).

It is worth exploring whether placing these genes into new genomic contexts could result in altered functions. Although transcriptome data in different species showed that specific gene transposition potentially inactivate these genes in different tissues (root, stem, leaf, flower, and ripe fruit) under normal growth conditions (Figure 6B), it is possible that they may exhibit novel features or have other regulatory functions, such as activation under adverse conditions. Further exploration and testing are needed to address these hypothese and clarify the mechanisms underlying the expansion and evolution of the SOT gene family in Rosaceae.

During our investigation of chromosomal location and synteny relations of genes, we observed a phenomenon that caught our attention: most paired genes from the same subfamily were distributed on two different chromosomes, and the sequence identity of pairs between two chromosomes was higher than that within the tandem cluster of the same chromosome. These results suggest that the ancestor genes of each SOT subfamily underwent differentiation events before whole genome duplication (WGD), If this were not the case, closely linked tandem gene pairs on the same chromosome would have exhibited high sequence identity and synteny.

To gain a better understanding of the evolution of SOT genes, we conducted a deep genealogy analysis and propose an evolutionary diagram that illustrates how one ancestral locus that predates the last common ancestor of Amygdaloideae gave rise to a large SOT gene clade with many subfamilies (Figure 7).

In conclusion, our study used a phylogenomic approach combined with synteny relationship analysis to provide insights into the early origin and evolution of SOT genes, as well as the complex gene duplication pattern of SOTs. The identification and evolutionary history of SOTs provide an intriguing example of how particular gene families evolve and expand during the evolution of species genomes.

## Materials and Methods

### Plant Genomes Collected

Sixty-one plant genomes were included in this study, with representation from various taxonomic groups including 34 rosids, 6 asterids, 7 early diverging eudicots, 10 monocots, 2 magnoliids, and 2 basal-angiosperms (*Euryale ferox* and *Amborella trichopoda*). For each genome, we retained only the primary transcript of protein sequences and the corresponding BED/GFF file information. The research method of SOT in 19 Rosaceae species was the same as that described above. Supplemental Table 1 and Supplemental Table 2 provide detailed information of genomes included in this study.

### PLT-targeting HMM Constructed and sequence identification

To identify the complete set of PLT sequences, we constructed a PLT-targeting Hidden Markov model (HMM) for monosaccharide transporters (MST). We obtained the HMM profile of the Sugar_tr domain (PF00083) from the Pfam database (http://pfam.xfam.org/), which was used to identify MST sequences.

Using HMMER3 (Finn et al., 2011), we searched and extracted all MST sequences with an E value below 0.001 from the genomes of eight representative species. We then selected 23 representative protein sequences of PLT and non-PLT from the MST gene family. To identify PLT-specific motifs, we conducted a motif analysis of the 46 sequences using the MEME online server (http://meme-suite.org/tools/meme), and the results are shown in Supplemental Figure 2.

We used the unique amino acids of PLT-specific motifs to construct the PLT-targeting HMM profile with the “hmmbuild” module of HMMER3. We then confirmed the reliability of the model by re-searching the original protein sequences of *Arabidopsis thaliana*, peach, and Dryas using the PLT-targeting HMM.

In the set of fifty-two protein sequences obtained above, we identified PLT protein sequences using the constructed model (PLT-targeting HMM). We conducted a search using hmmsearch program with a gathering threshold E-value < 1E-03 in HMMER3. We labeled PLT clade sequences based on the *Arabidopsis thaliana* sequences.

### Identification of SOT family members in Rosaceae plants

To identify SOT sequences and reveal the evolutionary relationship of SOT among the species of Rosaceae, we selected a total of 19 species of the Rosaceae, with *Arabidopsis*, grape (*Vitis vinifera*) and tomato (*Solanum lycopersicum*) as the outgroup. The protein sequences of reported SOT (Supplemental Table 3) were used as query sequences to retrieve SOT homologous sequences from the already available PLT protein dataset.

### Sequence alignment and phylogenetic analysis

Multiple alignments of protein sequences were performed using Muscle5 and MAFFT with the L-INS-I strategy (Katoh & Standley, 2013, Edgar, 2022). Poorly alignment positions and spurious sequences were removed using trimAl v1.2rev59 with the “-seqoverlap 50-resoverlap 0.5” (Salvador et al., 2009). Maximum-likelihood trees were constructed using IQtree v2.1.2 under the JTT+ R7 protein model with 1000 bootstrap replicates (Kalyaanamoorthy et al., 2017, Nguyen et al., 2015). Phylogenetic trees were visualized and plotted using iTOL (http://itol.embl.de) (Letunic & Bork, 2021).

### Syntenic block detection and synteny network constrction

We followed the method proposed by Zhao and Schranz (Zhao & Schranz, 2017) to calculate synteny blocks and construct synteny networks. All-to-all reciprocal whole-genome protein comparisons for 61 species was performed using Diamond v0.9.14 (Buchfink et al., 2021). All syntenic blocks between and within species genomes were detected by MCScanX with default parameters (Wang et al., 2012). Syntenic blocks associated with target genes were extracted from the entire synteny block database to obtain the final synteny network. Phylogenomic profiles were produced by counting the number of syntenic genes in each genome for each synteny cluster. The network was displayed and manipulated in Cytoscape v3.3.0 and Gephi v0.9.1 (Shannon et al., 2003, Bastian et al., 2009).

### Microsynteny analysis of SOT and flanking Genes

We performed microsynteny analysis for SOT gene-centric blocks (SOT genes as well as 5 upstream and downstream flanking genes) in five representative Rosaceae species, including apple (*Malus domestica*), loquat (*Eriobotrya japonica)*, bowman’s root (*Gillenia trifoliata*), peach (*Prunus persica*), and strawberry (*Fragaria vesca*). The pairwise synteny regions were searched by MCScanX (Python version) (Tang et al., 2015, Tang et al., 2008). Microsynteny analysis and visualization was generated using JCVI (https://github.com/tanghaibao/jcvi/wiki/Mcscan-(python-version). As the same time, the genes were plotted on chromosomes using MG2C (Jiangtao Chao 2021) and R v4.2.0.

### Gene expression profiles

We downloaded different RNA-seq samples from comprehensive repositories, and the detailed information of RNA-seq is displayed in Supplemental Table 4. Raw data were filtered using Fastp v0.20.1 with the “-l 50” option (Chen et al., 2018). Clean reads of sample were mapped to coding sequences (CDS) with Kallisto v.0.46.1 to obtain transcripts per million (TPM) gene expression values (Bray et al., 2016).

## References

Aprea E, Charles M, Endrizzi I, et al., 2017. Sweet taste in apple: the role of sorbitol, individual sugars, organic acids and volatile compounds. Sci Rep 7, 44950.

Bastian M, Heymann S, Jacomy M, 2009. Gephi: An Open Source Software for Exploring and Manipulating Networks. Proc. Int. AAAI Conf. Weblogs Soc. Media 8, 361–362.

Buchfink B, Reuter K, Drost HG, 2021 Sensitive protein alignments at tree-of-life scale using DIAMOND, Nature Methods 18, 366–368.

Capella-Gutiérrez S, Silla-Martínez JM, Gabaldón T, 2009. trimAl: a tool for automated alignment trimming in large-scale phylogenetic analyses. Bioinformatics 25, 1972–3.

Edgar RC, 2022. Muscle5: High-accuracy alignment ensembles enable unbiased assessments of sequence homology and phylogeny. Nat Commun 13, 6968.

Eirín-López JM, Rebordinos L, Rooney AP, Rozas J, 2012. The birth-and-death evolution of multigene families revisited. Genome Dyn 7, 170–96.

Fan RC, Peng CC, Xu YH, et al., 2009. Apple sucrose transporter SUT1 and sorbitol transporter SOT6 interact with cytochrome b5 to regulate their affinity for substrate sugars. Plant Physiol 150, 1880–901.

Gabaldon T, Koonin EV, 2013. Functional and evolutionary implications of gene orthology. Nat Rev Genet 14, 360–6.

Gao Z, Maurousset L, Lemoine R, Yoo SD, Van Nocker S, Loescher W, 2003. Cloning, expression, and characterization of sorbitol transporters from developing sour cherry fruit and leaf sink tissues. Plant Physiol 131, 1566–75.

Gu C, Wu RF, Yu CY, et al., 2021. Spatio-temporally expressed sorbitol transporters cooperatively regulate sorbitol accumulation in pear fruit. Plant Science 303.

Hanada K, Zou C, Lehti-Shiu MD, Shinozaki K, Shiu SH, 2008. Importance of lineage-specific expansion of plant tandem duplicates in the adaptive response to environmental stimuli. Plant Physiol 148, 993–1003.

Letunic I, Bork P, 2021. Interactive Tree Of Life (iTOL) v5: an online tool for phylogenetic tree display and annotation. Nucleic Acids Research 49, W293–w296.

Jia J, Xie Y, Cheng J, Kong C, Wang M, Gao L, Zhao F, Guo J, Wang K, Li G, Cui D, Hu T, Zhao G, Wang D, Ru Z, Zhang Y, 2021. Homology-mediated inter-chromosomal interactions in hexaploid wheat lead to specific subgenome territories following polyploidization and introgression. Genome Biol 22, 26.

Jiangtao Chao ZL, Yuhe Sun, Oluwaseun Olayemi Aluko, Xinru Wu, Qian Wang and Guanshan Liu, 2021. MG2C: a user-friendly online tool for drawing genetic maps. Molecular Horticulture 1, 4.

Johnson DA, Hill JP, Thomas MA, 2006. The monosaccharide transporter gene family in land plants is ancient and shows differential subfamily expression and expansion across lineages. BMC Evol Biol 6, 64.

Johnson DA, Thomas MA, 2007. The monosaccharide transporter gene family in Arabidopsis and rice: a history of duplications, adaptive evolution, and functional divergence. Mol Biol Evol 24, 2412–23.

Kalyaanamoorthy S, Minh BQ, Wong TKF, von Haeseler A, Jermiin LS, 2017. ModelFinder: fast model selection for accurate phylogenetic estimates. Nat Methods 14, 587–589.

Katoh K, Standley DM, 2013. MAFFT multiple sequence alignment software version 7: improvements in performance and usability. Mol Biol Evol 30, 772–80.

Kerstens MHL, Schranz ME, Bouwmeester K, 2020. Phylogenomic analysis of the APETALA2 transcription factor subfamily across angiosperms reveals both deep conservation and lineage-specific patterns. Plant Journal 103, 1516–24.

Klepek YS, Volke M, Konrad KR, et al., 2010. Arabidopsis thaliana POLYOL/ 1 and 2: fructose and xylitol/H+ symporters in pollen and young xylem cells. J Exp Bot 61, 537–50.

Kong W, Sun T, Zhang C, Qiang Y, Li Y, 2020. Micro-Evolution Analysis Reveals Diverged Patterns of Polyol Transporters in Seven Gramineae Crops. Front Genet 11, 565.

Lam-Tung Nguyen, Heiko A. Schmidt, Arndt von Haeseler, Bui Quang Minh, 2015. IQ-TREE: A Fast and Effective Stochastic Algorithm for Estimating Maximum-Likelihood Phylogenies. Molecular Biology and Evolution 32, 268–274

Li M, Feng F, Cheng L, 2012. Expression patterns of genes involved in sugar metabolism and accumulation during apple fruit development. PLoS One 7, e33055.

NL Bray, H Pimentel, P Melsted, L Pachter, 2016. Near optimal probabilistic RNA-seq quantification, Nature Biotechnology 34, 525–527.

Panchy N, Lehti-Shiu M, Shiu SH, 2016. Evolution of Gene Duplication in Plants. Plant Physiol 171, 2294–316.

Qiao X, Li Q, Yin H, et al., 2019. Gene duplication and evolution in recurring polyploidization-diploidization cycles in plants. Genome Biol 20, 38.

Qiao X, Yin H, Li L, et al., 2018. Different Modes of Gene Duplication Show Divergent Evolutionary Patterns and Contribute Differently to the Expansion of Gene Families Involved in Important Fruit Traits in Pear (Pyrus bretschneideri). Front Plant Sci 9, 161.

Reams AB, Roth JR, 2015. Mechanisms of gene duplication and amplification. Cold Spring Harb Perspect Biol 7.

Reinders A, Panshyshyn JA, Ward JM, 2005. Analysis of transport activity of Arabidopsis sugar alcohol permease homolog AtPLT5. J Biol Chem 280, 1594–602.

Robert D Finn, Jody Clements, Sean R. Eddy, 2011. HMMER web server: interactive sequence similarity searching. Nucleic Acids Research 39, W29–W37.

Robert J Schmitz, Erich Grotewold, Maike Stam, 2003. Cis-regulatory sequences in plants: Their importance, discovery, and future challenges. The Plant Cell 34, 718–741.

Shannon P, Markiel A, Ozier O, et al., 2003. Cytoscape: a software environment for integrated models of biomolecular interaction networks. Genome Res 13, 2498–504.

Shifu Chen, Yanqing Zhou, Yaru Chen, Jia Gu, 2018. fastp: an ultra-fast all-in-one FASTQ preprocessor. Bioinformatics 34, i884–i890

Slawinski L, Israel A, Paillot C, et al., 2021. Early Response to Dehydration Six-Like Transporter Family: Early Origin in Streptophytes and Evolution in Land Plants. Frontiers in Plant Science 12.

Stokes ME, Chattopadhyay A, Wilkins O, Nambara E, Campbell MM, 2013. Interplay between Sucrose and Folate Modulates Auxin Signaling in Arabidopsis. Plant Physiology 162, 1552–65.

Tang HB, Krishnakumar V, Li J, Zhang, X, 2015. jcvi: JCVI utility libraries: Zenodo. 10.5281/zenodo.31631.

Tang HB, Bowers JE, Wang XY, et al., 2008. Perspective - Synteny and collinearity in plant genomes. Science 5875, 486–488

Wang JP, Nayak S, Koch K, Ming R, 2013. Carbon partitioning in sugarcane (Saccharum species). Frontiers in Plant Science 4.

Wang L, Qi X, Yang Y, Zhang S, 2016. Molecular characterization and expression pattern of sorbitol transporter genePbSOT2in Pear (Pyrus bretschneideriRehd.) fruit. Canadian Journal of Plant Science 96, 128–37.

Wang Y, Tang H, Debarry JD et al., 2012. MCScanX: a toolkit for detection and evolutionary analysis of gene synteny and collinearity. Nucleic Acids Res 40.

Watari J, Kobae Y, Yamaki S, et al., 2004. Identification of sorbitol transporters expressed in the phloem of apple source leaves. Plant Cell Physiol 45, 1032–41.

Wu J, Wang Y, Xu J, et al., 2018. Diversification and independent domestication of Asian and European pears. Genome Biol 19, 77.

Yu C-Y, Cheng H-Y, Cheng R, et al., 2019. Expression analysis of sorbitol transporters in pear tissues reveals that PbSOT6/20 is associated with sorbitol accumulation in pear fruits. Scientia Horticulturae 243, 595–601.

Zhang HP, Wu JY, Tao ST, et al., 2014. Evidence for Apoplasmic Phloem Unloading in Pear Fruit. Plant Molecular Biology Reporter 32, 931–9.

Zhang K, Wang X, Cheng F, 2019. Plant Polyploidy: Origin, Evolution, and Its Influence on Crop Domestication. Horticultural Plant Journal 5, 231–9.

Zhang X, Li X, Zhao R, Zhou Y, Jiao Y, 2020. Evolutionary strategies drive a balance of the interacting gene products for the CBL and CIPK gene families. New Phytol 226, 1506–16.

Zhao T, Holmer R, De Bruijn S, Angenent GC, Van Den Burg HA, Schranz ME, 2017. Phylogenomic synteny network analysis of MADS-Box Transcription Factor Genes Reveals Lineage-Specific Transpositions, Ancient Tandem Duplications, and Deep Positional Conservation. Plant Cell 29, 1278–92.

Zhao T, Schranz ME, 2017. Network approaches for plant phylogenomic synteny analysis. Curr Opin Plant Biol 36, 129–34.

Zhao T, Schranz ME, 2019. Network-based microsynteny analysis identifies major differences and genomic outliers in mammalian and angiosperm genomes. Proc Natl Acad Sci U S A 116, 2165–74.

Zhu X, Tang C, Li Q, et al., 2021. Characterization of the pectin methylesterase inhibitor gene family in Rosaceae and role of PbrPMEI23/39/41 in methylesterified pectin distribution in pear pollen tube. Planta 253, 118.

